# ProLIF: a quantitative assay for investigating integrin cytoplasmic protein interactions and synergistic membrane effects on proteoliposomes

**DOI:** 10.1101/209262

**Authors:** Nicola De Franceschi, Mitro Miihkinen, Hellyeh Hamidi, Jonna Alanko, Anja Mai, Laura Picas, Daniel Lévy, Peter Mattjus, Benjamin T Goult, Bruno Goud, Johanna Ivaska

## Abstract

Integrin transmembrane heterodimeric receptors control a wide range of biological interactions by triggering the assembly of large multiprotein complexes at their cytoplasmic interface. A diverse set of methods have been used to investigate cytoplasmic interactions between integrins and intracellular proteins. These predominantly consist of peptide-based pull-downs and biochemical immuno-isolations from detergent-solubilized cell lysates. However, quantitative methods to probe integrin-protein interactions in a more biologically relevant context where the integrin is embedded within a lipid bilayer have been lacking. Here we describe a technique called ProLIF (<u>Pro</u>tein-<u>L</u>iposome <u>I</u>inenteractions by <u>F</u>low cytometry) to reconstitute recombinant integrin transmembrane domain (TMD) and cytoplasmic tail (CT) fragments on liposomes as individual α or β subunits or as αβ heterodimers and, using flow cytometry, to rapidly and quantitatively measure protein interactions with these membrane-embedded integrins. Importantly, the assay can analyse binding of fluorescent proteins directly from cell lysates without further purification steps. By combining integrins with membrane lipids to generate proteoliposomes, the effects of membrane composition such as PI(4,5)P_2_ presence on protein recruitment to the integrin CTs can be analyzed. ProLIF requires no specific instrumentation, apart from a standard flow cytometer and can be applied to measure a broad range of membrane-dependent protein-protein interactions with the potential for high-throughput/multiplex analyses.

## Introduction

Lipids provide an essential platform for protein interactions and biochemical reactions at biological membranes. Many techniques are available to assess protein-lipid binding and phosphoinositide (PI) specificity (Zhao and Lappalainen, 2012). Many of these assays and in particular those based on liposome generation - currently considered more representative of the *in cellulo* situation - need specialized equipment or employ complex protocols (e.g. surface plasmon resonance, isothermal titration calorimetry and lipid microarray) (Lemmon *et al.*, 1995; Ananthanarayanan *et al.*, 2003; Besenicar *et al.*, 2006; Wu *et al.*, 2012; Saliba *et al.*, 2014) that restrict their usage to specialized laboratories. Furthermore these approaches require high lipid/protein concentrations that prevent large and systematic analyses and/or remain merely qualitative. Recently, several microscopy-based methods have been developed (Ceccato *et al.*, 2016; Saliba *et al.*, 2014) that provide quantitative data on protein interactions with liposomes and have the potential for high-throughput analyses. Flow cytometry has also been employed to quantify binding of purified recombinant proteins to liposomes (Temmerman and Nickel, 2009). However, none of these methodologies have been designed to incorporate transmembrane proteins within the lipid bilayer.

It is estimated that transmembrane proteins constitute up to one third of the human proteome (Ahram *et al.*, 2006; Almen *et al.*, 2009) and are essential components of biological membranes, constituting approximately 50% of the membrane volume (Müller *et al.*, 2008). Transmembrane proteins regulate a plethora of essential cellular events, ranging from signal transduction to the flux of ions and metabolites across the membrane in response to a changing microenvironment. Due to their functions and accessibility, they represent more than 60% of drug targets (Arinaminpathy *et al.*, 2009). In spite of their importance, versatile methodologies to explore protein-protein interactions of transmembrane proteins within an experimentally controlled lipid microenvironment remain underdeveloped. Integrins, an essential family of heterodimeric transmembrane adhesion receptors, recruit and support the formation of cytoplasmic protein complexes, collectively known as the integrin adhesome, at the plasma membrane to generate the cell machinery responsible for cell adhesion and adhesion-induced signaling and migration (Winograd-Katz *et al.*, 2014).

Currently, molecular interactions between integrin and adhesome components are mainly studied by qualitative techniques such as pull-downs using synthetic peptides or soluble recombinant proteins mimicking the integrin cytoplasmic domains. Alternatively, endogenous integrins are immunoprecipitated in the presence of detergents. In all these approaches, an intact membrane is absent, even though several core adhesome proteins, such as talin, are known to bind acidic phospholipids. As a result, investigations into the joint requirement of integrin TMD-CT domains and acidic phospholipids in mediating protein recruitment to integrin tails have been, thus far, largely neglected.

Here, we describe a simple, sensitive and quantitative technique called ProLIF (<u>Pro</u>tein-<u>L</u>iposome <u>I</u>nteractions by <u>F</u>low cytometry) to simultaneously detect and quantify protein-protein and protein-lipid interactions in reconstituted proteoliposomes. We reconstituted “artificial integrins” into proteoliposomes and investigated talin binding, as it is the most studied protein interacting with both the integrin cytoplasmic tail and the plasma membrane in a phosphatidylinositol phosphate (PIP)-dependent manner (Calderwood *et al.*, 2013). We used this interaction to demonstrate the applicability of our method for probing integrin-cytoplasmic protein interactions in the context of a lipid bilayer of defined composition. We optimized ProLIF towards a mammalian expression system to circumvent the requirement for protein purification, preserve post-translational modifications, and to enable the presence of possible essential co-factors to provide a more realistic biological characterization of protein-protein binding.

## Results

### Generation of streptavidin-bead coupled liposomes for FACS detection

We first tested ProLIF by analyzing the coupling of bare liposomes, containing a small fraction of biotinylated-lipids, to streptavidin-coated carrier beads, according to steps 1, 3 and 4 outlined in the workflow in Fig.1a. Liposomes are produced by lipid solubilization in Triton X-100 and subsequent detergent removal by gradual addition of Bio-Beads^TM^ (Rigaud *et al.*, 1995). Although bare liposomes can also be produced using extrusion, giving control over the size of the resulting small unilamellar vesicles (SUVs) (Temmerman and Nickel, 2009), this technique does not allow for incorporation of transmembrane proteins. In contrast, detergent removal by Bio-Beads^TM^ is a robust method that has been used to reconstitute many functional transmembrane proteins (Mouro-Chanteloup *et al.*, 2010; Young *et al.*, 1997; Richard *et al.*, 1990; Moriyama *et al.*, 1984; Neves *et al.*, 2009; Lacapêre *et al.*, 2001; Geertsma *et al.*, 2008; Kolena, 1989; Nesper *et al.*, 2008; Smith and Morrissey, 2004) resulting in unilamellar vesicles (Rigaud *et al.*, 1995). Such vesicles are close to the detection limit of the scatter of laser light in FACS instruments (Temmerman and Nickel, 2009). In order to make these liposomes amenable to standard flow cytometry detection, we incorporated biotinylated lipids (2% of total lipid content) during liposome preparation to enable vesicle capture on Streptavidin Sepharose beads (SA)-beads that have an average diameter of 34 μm. The SA-beads are easily detected in a flow cytometer using forward scatter (FSC) and side scatter (SSC) plots (Fig.S1a). Upon addition of biotinylated liposomes, a distinct population of small objects appears (Fig.S1b); however, this population was gated out during the analysis. Importantly, addition of biotinylated liposomes did not appear to promote bead aggregation, as the FSC-A (Forward scatter area)/FSC-W (Forward scatter width) plot demonstrated a single population. To confirm that liposomes were captured by the SA-beads, we produced liposomes encapsulating Cy5 dye (Fig. 1b). A strong signal was detected by flow cytometry when the Cy5-encapsulated liposomes where captured on SA-beads. Importantly, interactions between SA-beads and Cy5-encapsulated biotinylated-liposomes could be effectively outcompeted by the addition of soluble biotin (Fig.1b), confirming specific biotin-mediated binding of liposomes to the carrier beads.

**Figure. 1.**
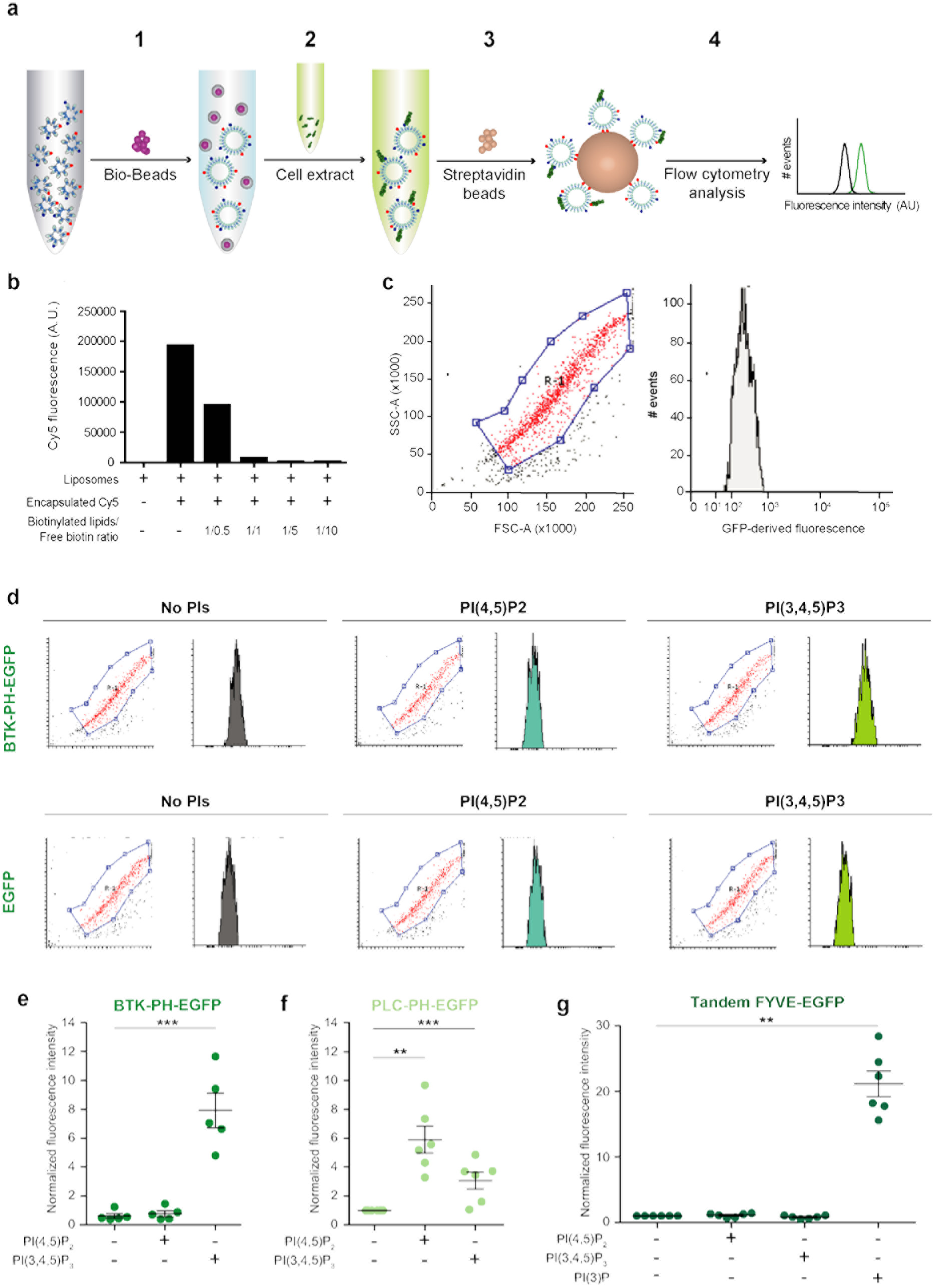
ProLIF is a flow-cytometry based assay for detection of specific protein-lipid interactions. **a**: Outline of ProLIF workflow. Step 1: Bio-Beads^TM^ are added to lipids solubilized in Triton X-100 to remove the detergent and obtain liposomes. Step 2: liposomes are incubated with membrane-free cell extract containing the EGFP-tagged protein of interest. Step 3: Sepharose streptavidin beads are added in order to capture the liposomes via interaction with biotinylated lipids present in the liposome membrane. Step 4: streptavidin beads are analyzed by flow cytometry (FACS). Red dots and blue dots represent biotinylated lipids and PIs, respectively. Green fragments represent EGFP-tagged proteins from the cell lysate. **b:** Biotinylated-lipid-containing liposomes were generated with and without encapsulated Cy5-dye, captured on SA-beads in the presence or absence of increasing amounts of free biotin and analyzed using FACS. Molar ratio between biotinylated lipids and soluble biotin added in each sample is indicated (n = 1). **c**: Scatter plot and fluorescence histograms from SA-beads alone incubated with cell lysate from EGFP transfected cells and analyzed by FACS. **d:** SA-beads coupled to biotinylated-lipid-containing liposomes, with the indicated PI content, were incubated with cell lysate from EGFP alone or BTK-PH-EGFP transfected cells (equal EGFP concentrations) and analyzed by FACS. Shown are representative dot blots, and size gating in FACS, and histograms depicting EGFP fluorescence intensity (FL1) profiles (note that the Axis labels are as in c). The red population in the scatter plot was gated for quantification. Data shown represent three individual experiments. **e**: Binding of BTK-PH-EGFP domain (from cell lysate as in d) to biotinylated-lipid-containing liposomes, with the indicated PI content, relative to control PI-free liposomes (n = 5, **p < 0.01, *** p < 0.001). **f**: Binding of EGFP-tagged PLC-PH domain (from cell lysate) to biotinylated-lipid-containing liposomes, with the indicated PI content, relative to control PI-free liposomes (n = 5, **p < 0.01, *** p < 0.001). **g**: Binding of tandem FYVE-EGFP domains (from cell lysate) to biotinylated-lipid-containing liposomes, with the indicated PI content, relative to PI-free liposomes (n = 6, **p < 0.01).

### Optimal detection of lipid interactions with proteins isolated from mammalian cell lysates

Protein purification can be time consuming and depending on the protein production source, critical post-translational modifications regulating protein binding to cell membrane components may be lacking. To overcome this limitation we tested the suitability of ProLIF to detect membrane interactions of phosphatidylinositide (PI)-binding proteins generated in human embryonic kidney cells (HEK293 cell line). Cells expressing EGFP-tagged PI-binding domains, known to interact with specific PIPs in membranes, were lysed in a detergent-free extraction buffer and fractions enriched in cytoplasmic proteins and devoid of transmembrane and membrane-associated molecules were isolated by ultracentrifugation (Fig.S1c). To overcome experimental variability due to changes in protein expression levels and to allow comparison between different experimental conditions, the fluorescence intensity of the cytoplasmic fractions were measured in relation to an external fluorescein standard and equalized before the binding assay.

Detergent-free cell lysates were subsequently incubated with liposomes followed by SA-beads and then liposome-bound SA-beads were analyzed by flow cytometry, according to the steps indicated in Fig.1a. All the cytometer settings (count rate, gates, voltages, trigger strategy) and the sample preparation conditions were kept constant for all samples. Beads were gated based on forward and side scattering and fluorescence intensity of the gated population was visualized using a histogram (fluorescence intensity vs. particle count) (Fig.1c,d).

SA-beads have a detectable level of auto-fluorescence (Fig. 1c), thus in each experiment a sample containing beads only was also included and the auto-fluorescence was subtracted from all samples. Thus, the specific fluorescence signal corresponding to EGFP-protein-bound liposomes was obtained. To determine the conditions providing the best signal to noise ratio, decreasing amounts of the phospholipase C-delta 1 (PLCδ1) pleckstrin homology (PH) domain (PLC-PH-EGFP), which binds preferentially to PI(4,5)P_2_ (Lemmon and Ferguson, 2000) were incubated with a constant amount of bare biotin-liposomes or PI(4,5)P_2_-containing biotin-liposomes, captured on SA-beads and analyzed by flow cytometry. The resulting titration data indicated that a concentration close to 8 nM provided a good compromise between achieving optimal signal/noise ratio and minimizing the amount of biological material needed for the experiment (Fig.S1d, see below the equation for calculating the protein concentration).

### Detecting specific protein-lipid interactions

Having established optimal experimental conditions to detect binding of fluorescently tagged proteins to liposomes, we next investigated whether ProLIF could be used to detect well-documented protein-lipid interactions in a reproducible manner. PH domains are broadly expressed in numerous cytoplasmic signalling proteins and are known to promote protein binding to specific lipids in the membrane. We first compared binding of EGFP alone or EGFP-tagged Bruton Tyrosine Kinase (BTK) PH domain (BTK-PH-EGFP) to various liposomes. Beads alone were used as a control for autofluorescence (as described above). In addition, bare liposomes (no PI) were compared to liposomes containing 2.5% PI(4,5)P_2_ or PI(3,4,5)P_3_. As shown in Fig. 1d, 1e; and S2a, binding of EGFP alone demonstrated background level binding with the signal intensity remaining similar in all liposome conditions. In contrast, BTK-PH-EGFP bound efficiently to PI(3,4,5)P_3_ liposomes, whereas binding to PI(4,5)P_2_ was very low, in line with the previously reported PI specificity for this PH-domain.

To explore the specificity of ProLIF further, we analyzed binding of two additional biologically distinct lipid-binding domains to liposomes. The PLCδ 1 PH-domain binds to PI(4,5)P_2_ serving as a specific tether that guides the protein to the plasma membrane (Garcia *et al.*, 1995). In contrast, the zinc-finger FYVE-domain, found in proteins such as the early endosomal antigen 1 (EEA1), binds phosphatidylinositol 3-phosphate (PI(3)P) specifically enriched on endosomal membranes, and fluorescently tagged fusions of tandem FYVE-domains (2xFYVE) serve as faithful reporters of PI(3)P enriched membranes in cells (Gillooly *et al.*, 2000; Stenmark *et al.*, 2002). Importantly, the PI specificity of both of these lipid-binding domains was recapitulated with ProLIF. We detected PLC-PH-EGFP binding specifically to liposomes containing 2.5% PI(4,5)P_2_ (Fig. 1f, S2b) and strong binding of a tandem FYVE zinc finger domain to PI(3)P (Fig. 1g).

### Quantitative analyses of protein-lipid interactions

To take the system a step further towards quantitative measurement of protein-lipid interactions we first devised a way to calculate the concentrations of the EGFP-tagged proteins in the input mammalian cell lysates by using an external fluorescein standard. Based on the measured lysate fluorescence, a mathematical equation (equation (2); eq.2) was derived (see methods) to calculate EGFP-tagged protein concentration as follows:

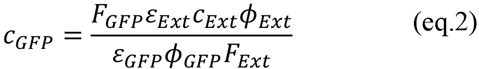

Where *C*_*Ext*_ and *C*_*GFP*_ are the concentrations of external standard (fluorescein) and the EGFP-tagged protein, *Φ*_*Ext*_ and *Φ*_*GFP*_ are the quantum yields of external standard and the EGFP-tagged protein and *ε*_*Ext*_ and *ε*_*GFP*_ are the extinction coefficients of external standard and the EGFP-tagged protein, respectively. To validate this equation, the fluorescence of a recombinant GFP protein of known concentration was measured at serial dilutions and a standard curve was generated. These experimentally derived fluorescence values were inputted into equation (2), together with variables and extinction coefficients from the fluorescein standard curve, and GFP concentrations were reverse calculated. Using this approach, a GFP standard curve closely matching the original experimental data was reproduced (Fig.2a). Mathematically derived standard curves for EGFP-tagged proteins were generated using predicted extinction coefficients (see methods) and quantum yields, and the fluorescence intensity of cell lysates expressing EGFP-tagged proteins of interest. Taking advantage of the calculated standard curve for the BTK-PH-EGFP, we incubated predetermined increasing concentrations of the protein with liposomes containing 2.5% PI(3,4,5)P_3_. As expected, and as demonstrated earlier with a similar approach for PH-PLCδ_1_ (Temmerman and Nickel, 2009), saturation of binding was achieved with increasing protein concentrations. Based on these data we calculated a K_D_ of 174 nM ± 15.2 (R^^2^ = 0.95) for BTK-PH-EGFP binding to PI(3,4,5)P_3_ (Fig.2b), which is within range of previously reported values (Kojima *et al.*, 1997; Ramesh *et al.*, 1997). However, in some cases the levels of expression of GFP-fused proteins in mammalian cell lysates may be incompatible for reaching saturation, limiting the applicability of this approach for some proteins in comparison to systems using purified recombinant proteins with liposomes (Temmerman and Nickel, 2009). With this limitation in mind, ProLIF is applicable for specific qualitative and quantitative analysis of biologically distinct protein-lipid interactions of proteins isolated from mammalian cell lysates.

**Figure 2.**
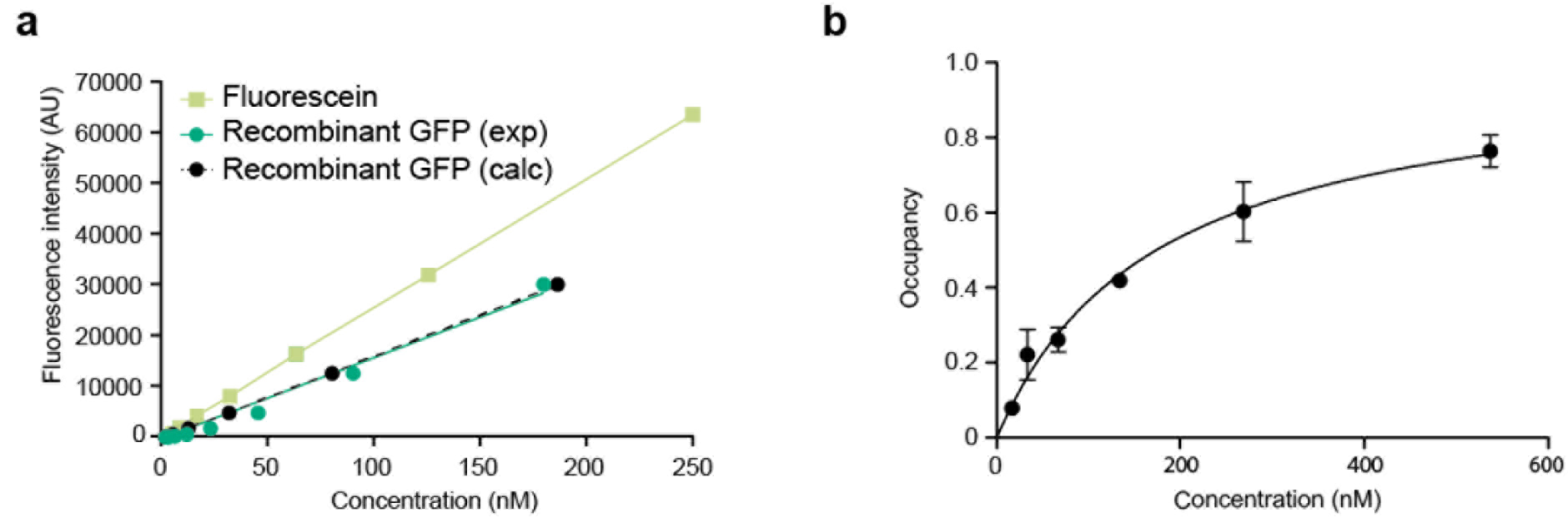
Quantitative measurements of protein-lipid interaction with ProLIF. **a**: Comparison of GFP and fluorescein standard curves. The fluorescence intensities of the indicated concentrations of fluorescein and recombinant GFP were determined experimentally (exp) and used to generate standard curves. The fluorescein standard curve was then used to calculate (calc) the theoretical GFP standard curve using equation (2). **b**: Titration curve of BTK-PH-EGFP binding to PI(3,4,5)P_3_-containing liposomes (n = 2). Cell lysates from BTK-PH-EGFP transfected cells were diluted to contain the indicated concentrations of the EGFP-tagged protein (calculated as in Fig. 2a using equation (2)) and incubated with the liposomes. Protein-liposome interactions were subsequently analysed by FACS as outlined by the workflow in Fig. 1a.

### Reconstituting integrin transmembrane-cytoplasmic domains on liposomes

To apply ProLIF to study transmembrane protein interactions, we chose integrins as model proteins. Integrin purification requires complex protocols that are not easy to scale up, precluding high-throughput application. For this reason, most of the studies involving purified full-length integrin are restricted to αIIbβ3, given the availability of platelets as a raw source. However, different integrin heterodimers can differ significantly in terms of physiological function and composition of their interactome (Rossier *et al.*, 2012). In order to overcome this limitation, we designed two artificial genes encoding the TMD and CT of the extracellular receptors α5 and β1 integrins and fused these to enhanced N-terminal Jun and Fos heterodimerization modules (cJun[R]-FosW[E]) (Worrall and Mason, 2011), respectively (Fig.3a) to promote α5 and β1 integrin pairing (integrins exist as heterodimers on the plasma membrane) in the same orientation. Such modular organization allows the study of different integrin heterodimers by simply modifying the TMD and cytoplasmic domains. Both Jun-α5 and Fos-β1 integrin chimeras could be purified from membrane fractions when expressed in *E. coli* by taking advantage of their purification tags, maltose binding protein (MBP) and glutathione S-transferase (GST), respectively (Fig.S3a,b). When analyzed by SDS-PAGE, both Jun-α5 (MW: 52.8 kDa) and Fos-β1 (MW: 40.7 kDa) protein bands, recognized by specific antibodies raised against the α5 and β1 integrin cytoplasmic domains, appeared at the correct size (Fig.3b). Moreover, Jun-α5 and Fos-β1 integrins were able to heterodimerize as demonstrated by reciprocal co-immunoprecipitation (co-IP) assays with antibodies against either the α5 or β1 integrin cytoplasmic domains (Fig.3b,c).

**Figure 3.**
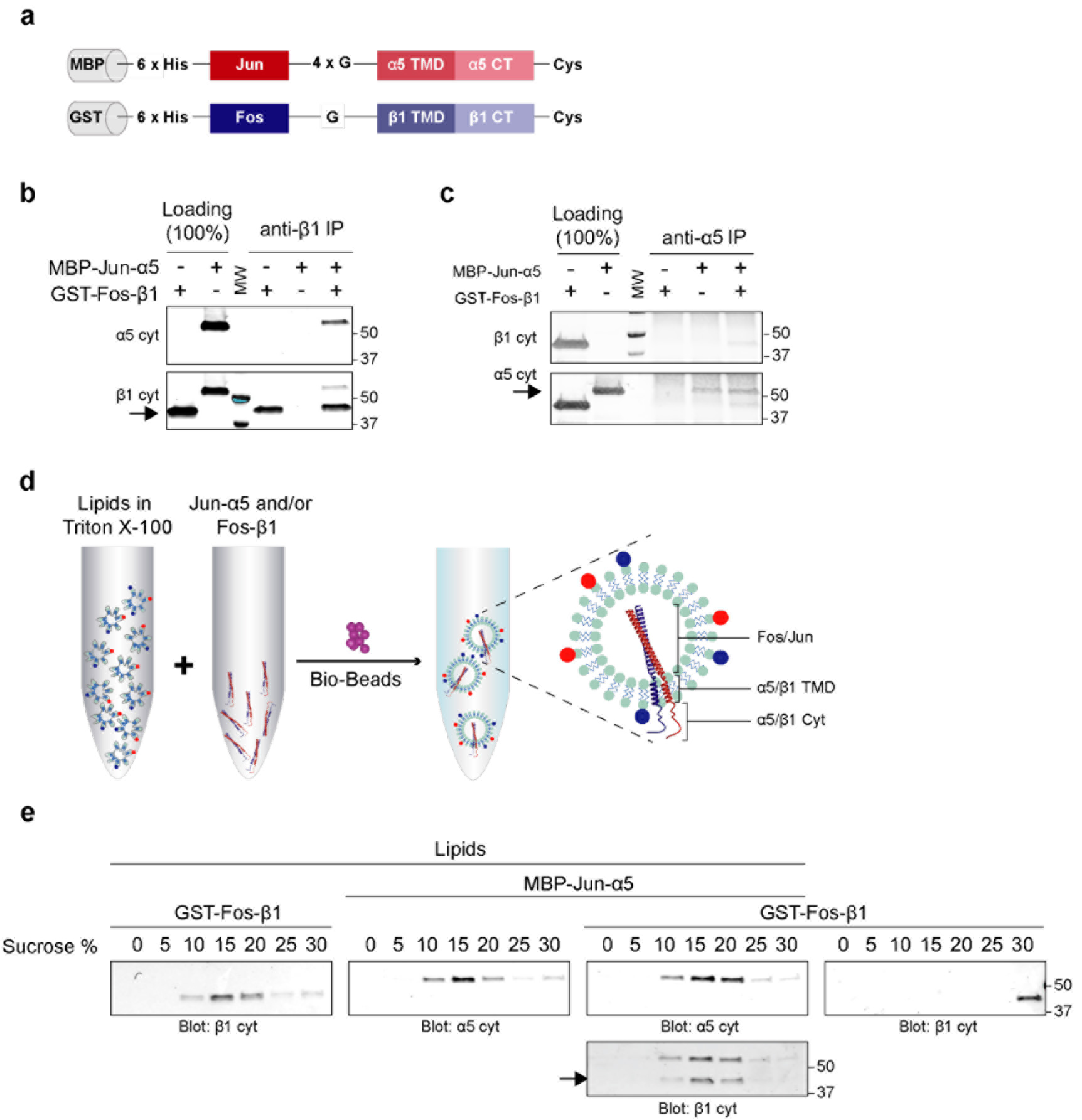
Reconstituting integrin transmembrane-cytoplasmic domains on liposomes. **a**: Domain architecture of MBP-Jun-α5 and GST-Fos-β1 constructs. G: glycine: TMD: transmembrane domain; CT: cytoplasmic domain; Cys: cysteine. **b:** The indicated purified recombinant proteins alone or in combination were subjected to immunoprecipitation with an anti-β integrin antibody directed against the β cytoplasmic domain: MBP-Jun-α5 co-immunoprecipitated with GST-Fos-β1. Filters were probed with rabbit anti-α5 integrin cytoplasmic domain antibody and then reprobed with rabbit anti-β1 integrin cytoplasmic domain antibody. Arrow indicates the β-integrin chimera band. Representative blot is shown. (n = 2 independent experiments). **c**: The indicated purified recombinant proteins alone or in combination were subjected to immunoprecipitation with an anti α5 integrin antibody directed against the α5 cytoplasmic domain: GST-Fos-β1 co-immunoprecipitated with MBP-Jun-ζ5. Filters were probed with rabbit anti-β1 integrin cytoplasmic domain antibody and then reprobed with anti-α5 integrin cytoplasmic domain antibody. Arrow indicates the ζ5-integrin chimera band. Representative blot is shown. (n = 2 independent experiments). **d**: Schematic of MBP-Jun-α5 and GST-Fos-β1 integrin incorporation in proteoliposomes. **e**: Gradient flotation assay showing reconstitution of GST-Fos-β1, MBP-Jun-α5 and the β1/α5 heterodimer in liposomes. Purified recombinant GST-Fos-β1 and MBP-Jun-α5 were incorporated either alone or in combination into liposomes as depicted in 3d. The resulting proteoliposomes were analyzed using a flotation assay in sucrose gradient. Liposome-incorporated-proteins float up the gradient (10-20% sucrose fractions), whereas in the absence of liposomes the protein alone remains in the bottom (30% sucrose) fraction (GST-Fos-β1 in the most right hand panel). The protein:lipid molar ratio is 1:3500 for both MBP-Jun-α5 and GST-Fos-β1. Arrow indicates the β1-integrin chimera in the reprobed filter.

Next, we reconstituted the Jun-α5 and/or Fos-β1 integrin chimeras in liposomes using the same protocol described above. The purified proteins, solubilized in mild detergent (see methods), were added to the Triton X-100 solubilized lipids, and incorporated into the lipid bilayer during detergent removal by Bio-Beads^TM^ (Fig. 3d). In this system, we lack the means to restrict the orientation of the fusion proteins on the liposomes resulting in approximately 50% of the reconstituted proteins having their cytoplasmic tails facing outwards. Given the strong affinity of the Jun-Fos dimer, in heterodimer-containing liposomes both α- and β-integrin tails are also expected to face the same way resulting in 50% of dimers having the correct orientation. To verify whether the purified proteins were indeed being incorporated into liposomes, we performed a sucrose gradient flotation assay. In the presence of liposomes, the integrin chimeras, as single entities or as components of a heterodimer, were retrieved from the upper sucrose fractions indicating association between the integrin proteins and the lipid bilayer (Fig.3e). In contrast, in the absence of lipids, protein aggregation was observed, and Fos-β1 was present in the lowest fraction (Fig.3e). Importantly, by using the Bio-Bead reconstitution method all protein is incorporated into liposomes, which makes a subsequent purification step (Ye *et al.*, 2010) unnecessary and helps to streamline the protocol.

### Integrin β1-cytoplasmic tail and PIPs synergize to recruit talin-head to liposomes

The integrin cytoplasmic domains have no enzymatic activity and function by recruiting, and binding to, cytoplasmic adaptors and signaling proteins that link the receptor to the actin cytoskeleton (Bouvard *et al.*, 2013). Talin is a classical integrin activator and one of the first proteins recruited to integrin heterodimers at the plasma membrane. The talin FERM domain binds directly to β integrin subunits, an event that is linked to separation of the α- and β-integrin tails and the subsequent activated conformation of the receptor and recruitment of other proteins. Talin also contains a PI binding surface within its FERM domain (Elliott *et al.*, 2010) and as such offers an excellent candidate for validating the ProLIF system. Using a concentration of EGFP-tagged talin FERM domain (3 nM) that was determined to provide a good signal/noise ratio (Fig.S3c), we observed significant talin binding to liposomes containing the Fos-β1 integrin protein (Fig.4a,b). As expected, talin did not bind to liposomes containing the Jun-α5 integrin subunit alone. Importantly, none of the conditions caused bead aggregation, as only a single main population was apparent in the FSC-A/FSC-W plots (Fig.S4). Interestingly, talin binding to the Fos-β1 integrin protein was completely lost when the β-integrin tail was embedded as part of the integrin heterodimer (Jun-α5-Fos-β1) within the liposome (Fig.4b), suggesting that this construct may represent a “tails-together” conformation of the integrin cytoplasmic face. This inhibitory effect was not due to membrane overcrowding, as reducing the transmembrane protein:lipid molar ratio by 50% (1:7000 instead of 1:3500) preserved the binding pattern (Fig.S5a). Talin FERM interaction with liposomes was modestly, but significantly, increased when PI(4,5)P2 was included in the liposomes, in line with the affinity of the talin FERM domain for plasma membrane acidic phospholipids (Calderwood *et al.*, 2013). Notably, the presence of PI(4,5)P_2_ and Fos-β1 integrin in the same liposomes substantially enhanced talin binding far beyond levels observed for each individual component, suggesting an additive and possibly synergistic binding effect, revealed by the ability of the ProLIF system to incorporate membrane-embedded integrins and membrane lipids in the same binding assay. In PI(4,5)P_2_ containing vesicles, talin FERM binding was reduced when both Fos-β1 and Jun-α5 were present (Fig.4b). Binding of talin FERM domain to Jun-α5 and PI(4,5)P_2_ was similar to conditions containing PI(4,5)P_2_ alone, suggesting that the talin FERM-PI(4,5)P_2_ interaction is preserved despite loss of interaction with the β1-integrin receptor (Fig.4b). Incubation with an excess of soluble biotin, which outcompetes liposome binding to the beads, resulted in the complete loss of the fluorescence signal (Fig.S5b), serving as an important control and confirming that the signal is only due to binding events occurring at the membrane rather than unspecific binding to the beads.

**Figure 4.**
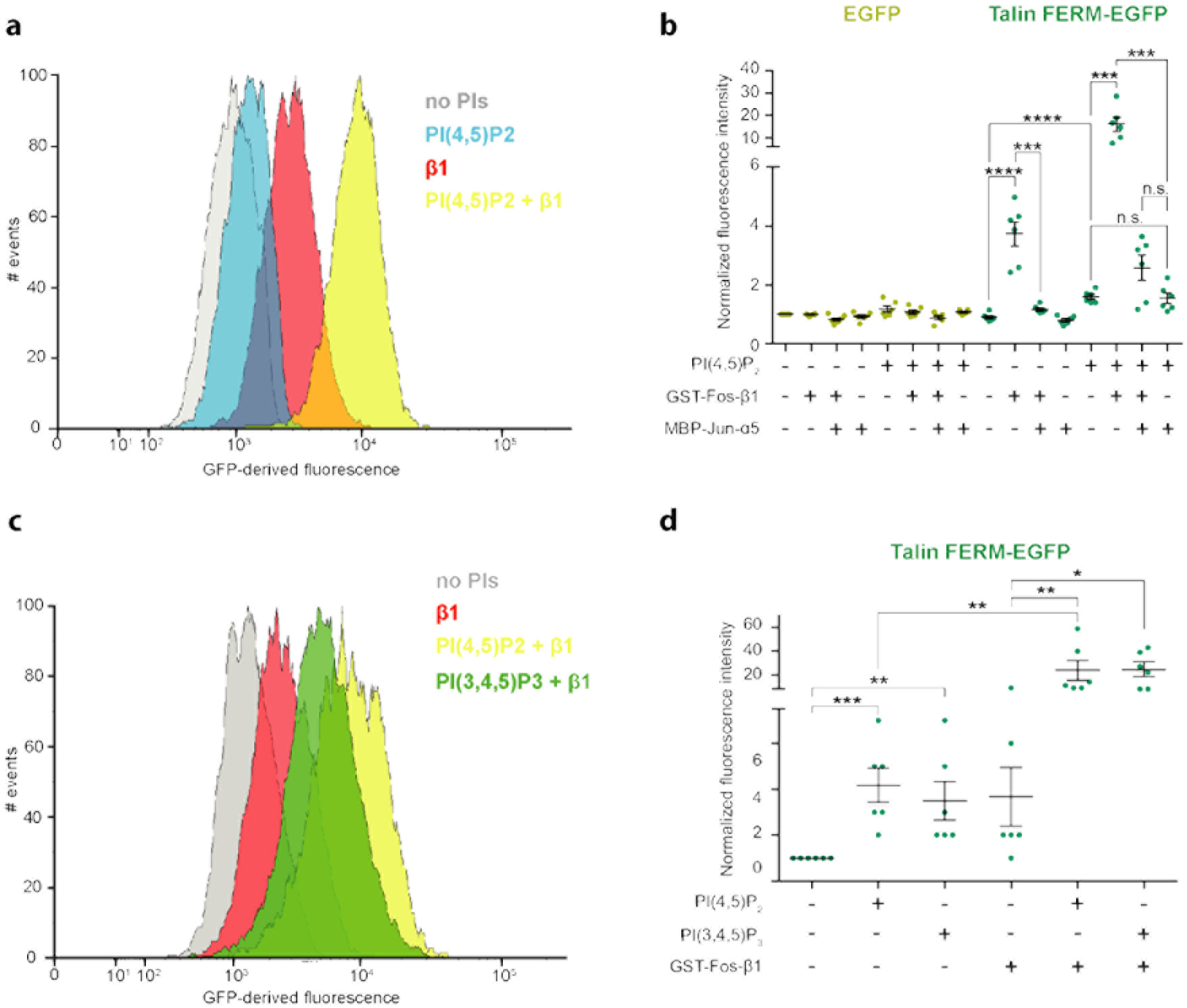
PIP2 and PIP3 synergize with integrin β1-tail to support talin-head recruitment. **a**: Example fluorescence intensity histograms of Talin FERM-EGFP (from lysates of transfected cells) binding to biotinylated-lipid-containing proteoliposomes, with PI(4,5)P_2_ and GST-Fos-β1 integrin as indicated. Gray, lipids control (no PI); blue, PI(4,5)P_2_; red, GST-Fos-β1; yellow, PI(4,5)P2 + GST-Fos-β1. **b**: Quantification of binding of Talin FERM-EGFP and EGFP control cell lysates at equimolar concentration to proteoliposomes with the indicated PI and integrin content (n = 4 independent experiments n.s. not significant, ***p < 0.001, ****p < 0.0001). **c**: Example fluorescence intensity histograms of Talin FERM-EGFP (from lysates of transfected cells) binding to biotinylated-lipid-containing proteoliposomes, with PI(4,5)P_2_, PI(3,4,5)P_3_ and GST-Fos-β1 integrin as indicated. Gray, lipids control (no PI); red, GST-Fos-β1; yellow, PI(4,5)P_2_ + GST-Fos-β1; green, PI(3,4,5)P_3_ + GST-Fos-β1. **d**: Quantification of binding of Talin FERM-EGFP to proteoliposomes with the indicated PI and integrin content. (n = 6 independent experiments *p < 0.05, **p < 0.01, ***p < 0.001, ****p < 0.0001).

With ProLIF, we could also observe talin binding to PI(3,4,5)P_3_ alone and detected a substantial enhancement of talin binding to PI(3,4,5)P_3_ and Fos-β1 integrin-containing liposomes that was equivalent to PI(4,5)P_2_ and Fos-β1 integrin-containing liposomes (Fig.4c,d). The ability of talin to tether to the β1-integrin cytoplasmic tail in conjunction with both PI(4,5)P_2_ and PI(3,4,5)P_3_ has not been carefully studied before and may be linked to interesting biological functions warranting further investigation in the future.

### Discussion

We demonstrate here that ProLIF is a sensitive, versatile and quantitative system to study protein interactions at the cytoplasmic interface of transmembrane proteins, taking into account the individual or synergistic contribution of protein-protein and protein-membrane lipid interactions.

The benefits and sensitivity of ProLIF are particularly exemplified with the integrin chimeras. Many individual protein-protein interactions in the integrin adhesome are characteristically of low affinity and much of the biology is based on synergistic binding events, clustering and multivalent interactions. Thus, studying the integrin cytoplasmic interactions with biochemical assays such as pull-downs with integrin-tail peptides in detergent can be challenging and do not represent the situation in cells. This is highlighted by the ProLIF data, which demonstrates that the talin-β1 integrin interaction is strongly enhanced by the presence of specific PI species. Thus, it is important to investigate how protein-protein interactions are regulated in the context of changing membrane lipid composition, an aspect that is potentially underestimated in the current integrin cell adhesion literature. Indeed, a number of lipid-binding domains have been identified and characterized (Lemmon, 2008) and the domain architecture of many proteins, including trafficking proteins, kinases and scaffold proteins, combines lipid- and protein-binding modules (Cullen, 2008; Pearce *et al.*, 2010). Thus, the synergistic effect observed for talin is likely to be a widespread phenomenon that could be addressed using ProLIF.

The mammalian expression system, optimized for ProLIF, also adds novelty over other methods available for monitoring protein-lipid binding as it supports posttranslational modifications of the soluble protein and the formation of protein complexes within cells. These events could be manipulated by biological reagents to gain further insight into mechanisms regulating protein binding to membrane components.

We believe that the simple strategy for lipid and protein reconstitution in liposomes and the use of a flow cytometer makes ProLIF a powerful, yet amenable tool for the quantitative detection of binding events on membranes, which can be applied to other transmembrane proteins. Moreover, ProLIF can be further developed into multiplexed assays by taking advantage of the palette of fluorescent tags available.

## Methods

**Plasmids and constructs**. The Jun-α5 integrin artificial gene (human alpha 5 amino acids 989-1049) was synthesized by DNA2.0 in pD441-HMBP. The Fos-β1 integrin (human beta 1 amino acids 725 to 798) artificial gene was synthesized by DNA2.0 and cloned in the pGEX-4T vector using EcoRI and BamHI cloning sites. A polyglycine linker was inserted between the Jun/Fos dimerization motifs and the integrin transmembrane domains. Insertion of the 6 x His Tag was performed by QuikChange II Site-Directed Mutagenesis Kit (Agilent Technologies). All constructs were fully sequenced prior to use.

The sequence of the Fos-β1 integrin chimera is as follows:GGATCCGGCGGCGGCGCGAGCCTGGATGAACTGGAAGCGGAAATTGAACAGCTGGAAGAAGAAAACTATGCGCTGGAAAAAGAAATTGAAGATCTGGAAAAAGAACTGGAAAAACTGGGCGCGCCGGGCACCGGCCCGGATATTATTCCGATTGTGGCGGGCGTGGTGGCGGGCATTGTGCTGATTGGCCTGGCGCTGCTGCTGATTTGGAAACTGCTGATGATTATTCATGATCGTCGTGAATTTGCGAAATTTGAAAAAGAAAAAATGAACGCGAAATGGGATACCGGCGAAAACCCGATTTATAAAAGCGCGGTGACCACCGTGGTGAACCCGAAATATGAAGGCAAATAATAAGAATTC

The sequence of the Jun-α5 integrin chimera is as follows:ATGGGATCCCATCATCATCATCATCATGGCGGCGGCGCGAGCATTGCGCGTCTGCGTGAA CGTGTGAAAACCCTGCGTGCGCGTAACTATGAACTGCGTAGCCGTGCGAACATGCTGCGTGAACGTGTGGCGCAGCTGGGCGCGCCGGGCGGCGGCGGCGGCACCAAAGCGGAAGGCAGCTATGGCGTGCCGCTGTGGATTATTATTCTGGCGATTCTGTTTGGCCTGCTGCTGCTGGGGCGATGGAAAAAGCGCAGCTGAAACCGCCGGCGACCAGCGATGCGTGCTAATAAGAATTC

Plasmids encoding BTK-PH-EGFP and PLC(δ1)-PH-EGFP were kind gifts from Matthias Wymann (University of Basel, Switzerland). The EGFP-tagged tandem FYVE was a gift from Harald Stenmark (Oslo University Hospital) and has been previously described (Gillooly *et al.*, 2000). The Talin FERM-EGFP (mouse talin1 residues 1-433) construct was made by the PROTEX facility at the University of Leicester, UK.

**Cells, antibodies, lipids and reagents**. HEK 293 cells (ATCC) were grown in Dulbecco’s modified Eagle’s medium (DMEM) with high glucose (4500 mg/ml) (Sigma-Aldrich) supplemented with 1% L-glutamine (Gibco), 10% fetal bovine serum (FBS; Sigma-Aldrich), and 1% penicillin-streptomycin (Sigma-Aldrich). The following antibodies were used: anti-integrin β1 Ab183666 (Abcam) and anti-integrin α5 AB 1949 (Millipore) for immunoblotting and immunoprecipitation. The following lipids were used: L-α-phosphatidylcholine (EggPC, 840051P); L-α-phosphatidic acid (EggPA, 840101P); 5-cholestene-3α,20α-diol (20α-hydroxycholesterol, 700156); L-α-phosphatidylinositol-4,5-bisphosphate (Brain PI(4,5)P2, 840046X); 1-stearoyl-2-arachidonoyl-*sn*-glycero-3-phospho-(1'-myo-inositol-3',4',5'-trisphosphate) (18:0-20:4 PI(3,4,5)P3, 850166P); 1,2-dioleoyl-*sn*-glycero-3-phospho-(1'-myo-inositol-3'-phosphate) (18:1 PI(3)P, 850150P); 1-oleoyl-2-(12-biotinyl(aminododecanoyl))-*sn*-glycero-3-phosphoethanolamine (18:1-12:0 Biotin PE, 860562P). All lipids were purchased from Avanti Polar Lipids. Recombinant GFP protein was purchased from Thermo Fischer Scientific. Streptavidin Sepharose High Performance beads (17-5113-01) were purchased from GE Healthcare.

**Membrane protein purification**. The Rosetta strain (Merck) of competent cells was used for Jun-α5 and Fos-β1 protein expression. Briefly, bacteria were transformed with the respective DNA according to manufacturer’s instructions and positive clones were selected on agar plates containing 100 μg/ml ampicillin and 33 μg/ml chloramphenicol (both from Sigma). Transformed bacteria were then grown in LB broth containing ampicillin and chloramphenicol until OD600 = 0.6 at which point protein expression was induced by addition of 0.5 mM IPTG (Sigma) for 5 hr at 25^o^C. Bacteria were pelleted, transferred to a falcon tube and flash-frozen in liquid N_2_. Cells were resuspended in 50 mM Tris pH 7.5, 150 mM NaCl, 600 μM TCEP (Tris(2-carboxyethyl)phosphine hydrochloride, Sigma), 500 μM PMSF (Sigma), 2 mM AEBSF (4-(2-Aminoethyl)benzenesulfonyl fluoride hydrochloride, Sigma), 0.1 mg/ml DNase (Roche), protease inhibitor (Roche), 1 mM mercaptoethanol (Sigma), 5 mM MgCl2 and lysozyme (Sigma) and disrupted using a cell disruptor. Cell lysates were clarified at 15000 rpm using a JA 25/50 rotor for 20 min at +4°C and resulting supernatants further centrifuged at 48000 rpm in a Ti50.2 rotor for 1 hr to pellet membranes. The membrane pellet was resuspended in 50 mM Tris pH 7.5, 150 mM NaCl, 600 μM TCEP, 500 μM PMSF, 1 mM AEBSF and homogenized in a Teflon homogenizer and after addition of sucrose (300 mM) samples were flash-frozen in liquid N_2_. Membrane suspensions were thawed, incubated with n-Dodecyl-β-D-Maltoside (DDM) (Anatrace) at a 5:1 (w:w) ratio for 2 hr at +4°C with agitation and centrifuged at 45000 rpm in a Ti50.2 rotor for 50 min at +4°C. Supernatants were incubated with Ni^2^+ sepharose beads (GE healthcare) for 2 hr at +4°C. Beads were washed with 50 mM Tris pH 7.5, 150 mM NaCl, 600 μM TCEP, 1 mM AEBSF + 0.5% DDM, followed by a second wash with 50 mM Tris pH 7.5, 150 mM NaCl, 600 μM TCEP, 1mM AEBSF and either 0.05% DDM (for Fos-β1) or 0.1% DDM (for Jun-α5). Proteins were eluted with 50 mM Tris pH 7.5, 150 mM NaCl, 1mM AEBSF, 0.05% DDM + 250 mM imidazole. Eluted proteins were incubated with either glutathione Sepharose beads (purification of GST-tagged Fos-b1; GE Healthcare) or dextrin sepharose beads (purification of MBP-tagged Jun-α5; GE Healthcare) for 60 min at +4°C. Beads were washed with 50 mM Tris pH 7.5, 150 mM NaCl, 600 μM TCEP, 1 mM AEBSF and either 0.05% DDM (for Fos-β1) or 0.1% DDM (for Jun-α5). Proteins were eluted with 50 mM Tris pH 7.5, 150 mM NaCl, 1 mM AEBSF, 0.05% DDM and either 30 mM glutathione (for Fos-β) or 20 mM maltose (for Jun-α5) and flash frozen in 10% glycerol in liquid N_2_ and stored at-8°C. Approximately 1 mg of protein/L of bacterial culture was purified using this technique.

**Bio-Beads^TM^ preparation and dosing**. Bio-Beads^TM^ (Bio-Rad) were sifted to exclude small beads and subsequently washed three times with methanol and five times with dH_2_O. Beads were left to sediment and during liposome preparation (see below) added in volumes of 15 μl (reproducibly corresponds to 3 mg of beads), collected from the bottom of the tube using a cut tip.

**Liposome and proteoliposome reconstitution**. The control lipid mix used throughout the study, unless otherwise indicated, was composed of 73% (w/w) Egg-PC, 10% (w/w) Egg-PA, 15% (w/w) cholesterol and 2% (w/w) biotinylated-lipids. Where indicated PIs were included at the expense of Egg-PA to preserve the percentage of negatively charged lipids at 10%.

In the case of BTK-PH-EGFP KD fitting and BTK-PH-EGFP example histograms in Fig. 1d the liposome composition used was 80.5% (w/w) POPC (synthetic substitute for Egg-PC; 850457P, Avanti Polar Lipids) lipids, 15% (w/w) cholesterol and 2% (w/w) biotinylated lipids + 2.5% (w/w) PI(3,4,5)P_3_.

The lipids solubilized in organic solvent were mixed and dried under a N_2_ stream, vacuum-dried for at least 20 min, resuspended in dH_2_O at 10 mg/ml and vortexed. The resulting liposomes were aliquoted in single-use aliquots and stored at –20°C. For each liposome/proteoliposome reconstitution, 400 μg of total lipids were solubilized in Triton X-100 (Triton X-100:lipid ratio of 2.5 w/w) in a total volume of 400 μl of reconstitution buffer (50 mM Tris pH 7.0, 150 mM NaCl and 600 μM TCEP) at RT with constant stirring until the solution became clear indicating total lipid solubilisation. Solubilized lipids were cooled to +4°C and 1 mM EDTA, 5 mM AEBSF, GST-Fos-β1 and/or MBP-Jun-α5 were added to the solution and stirred at +4°C for 15 min. Prewashed Bio-Beads^TM^ (total 48 mg) were gradually added to the solution at +4°C and stirred according to the following scheme:

**Table I.**

| **Bio-Beads™ aliquot (mg)** | **Incubation time at +4°C with**<br>**agitation** |
| --- | --- |
| **3** | 90 min |
| **3** | 90 min |
| **12** | Overnight |
| **30** | 120 min |

**Cell transfection**. HEK 293 cells were seeded at a density of 25-35% confluence and transfected the next day at 50-70% confluence according to the following protocol for a 10 cm dish. The plasmid of interest (12 μg) was mixed with 250 μ1 of Opti-MEM (Gibco) for 5 min at RT. A premix of PEI 25K transfection reagent solution (Polysciences Inc) (30 μ1 incubated with 250 μ1 of Opti-MEM for 5 min at RT) was then added and incubated for a further 10 min at RT. The transfection solution was placed on top of the 5 ml of culture medium present in the cell culture dish. Cells were harvested after overnight incubation.

**Isolation of detergent-free cell lysate**. Cells were washed twice with PBS and scraped in 400 μl of detergent-free lysis buffer (10 mM Tris pH 7.5, 250 mM sucrose, 1 mM EDTA, 1 mM MgOAc, 20 μM ATP + Complete protease and PhosSTOP phosphatase inhibitor tablets, Roche) at +4°C. Cell extracts were passed through a syringe needle (0.5 mm) five times, sonicated at +4°C for 5 min and ultracentrifugated at 100000 x g for 1 hr at +4°C. The resulting supernatant, depleted of membrane and transmembrane fractions, was used for the experiment.

**Co-immunoprecipitation**. An equimolar mixture of Fos-β and Jun-α5 were subjected to immunoprecipitation using 1 μg of the indicated antibodies at +4°C for 2 hr in 50 mM Tris pH 7.5, 150 mM NaCl, 600 μM TCEP + 0.1% DDM. Immunoprecipitated complexes were isolated on protein-G beads (GE Healthcare) for 2 hr at +4°C. Beads were then washed once with the same buffer and suspended into loading buffer. Samples were separated by SDS-PAGE and analyzed using Western blotting.

**Flotation assay**. Equimolar amounts of Fos-β and Jun-α5 were reconstituted in liposomes, mixed in 1:1 ratio with a 60% sucrose solution and added to the bottom of an ultracentrifuge tube. Decreasing concentrations of sucrose were progressively layered on top to form a gradient and the sample was centrifuged overnight at 20000 x g. Fractions were retrieved and analyzed by SDS-PAGE.

**Calculation of EGFP concentration within cell lysates**. The fluorescence intensity of serial dilutions of fluorescein (1 nM – 256 nM in dH_2_O) was measured using the BioTek Synergy H1 hybrid reader to obtain a standard curve. The fluorescence intensity of cell lysates was measured in relation to this standard curve and EGFP-tagged protein concentration calculated using equation (2). Fluorescein quantum yield in dH_2_O (ϕ_Ext_) and extinction coefficient (ɛ_Ext_) are 0.76 and 76900 M^−1^cm^−1^, respectively (Song *et al.*, 2000; Zhang *et al.*, 2014); GFP quantum yield (ϕ_gep_) and extinction coefficient (ɛ_gep_) are 0.53 and 70000 M^1^cm^1^ (for dimeric GFP) (Thermo Scientific), respectively. For EGFP-tagged proteins, extinction coefficients were calculated using the ExPASy ProtParam tool at http://web.expasy.org/protparam/ to obtain the predicted coefficient for each EGFP-tagged construct.

The fluorescence intensity or number of excited molecules during passage of light through a sample can be derived from the Beer-Lambert law (equation 1; eq.1):

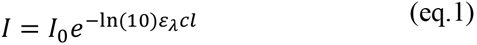

where (*I*) corresponds to the transmitted light through the sample, (*I*_0_) is the incident radiation, (*ɛ*_*λ*_) is the extinction coefficient at the excited wavelength (*λ*), (*c*) is the concentration, and (*l*) is the light path length. For low absorbance values, this can be expanded to:

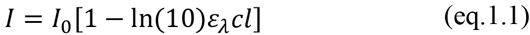

The emission intensity (*F*_*λ*_) for one type of molecule at a given wavelength is a function of the quantum yield (*ϕ*_*F*_), the fraction of emission that occurs at that wavelength (*f*_λ_), and the fraction of the radiation that is actually collected by the detector (*j*):

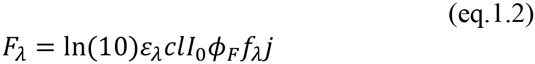

Solving this equation for the concentration of our EGFP-labelled molecule we obtain the following expression (sub-indices indicate the sample):

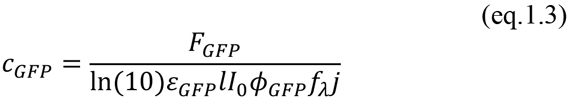

Now using the calibration curve obtained with external standard we can obtain the incident radiation
(I_0_):

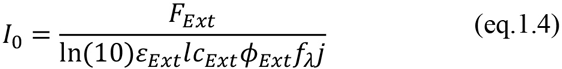

that when combined with the previous equation results in equation (2):

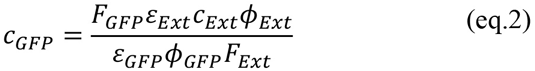

where the ratio 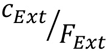is the inverse of the slope in the linear fit of *F*_*Ext*_ as a function of *c*_*Ext*_ in the calibration curve.

**Flow cytometry-based binding assay**. The fluorescence of cell lysate (excitation/emission, 485/528) was measured in relation to a fluorescein titration curve in dH_2_O using the BioTek Synergy H1 hybrid reader. Equation (2) was applied to calculate the actual EGFP-tagged protein concentration. The concentration of cell lysate was adjusted by dilution in detergent-free lysis buffer. 150 μ1 Cell lysate was transferred to an Eppendorf tube and incubated with 90 μ1 of reconstitution buffer (50 mM Tris pH 7.0, 150 mM NaCl and 600 μM TCEP) and 60 μl for 4 h with constant stirring at +4°C.

Samples were then incubated with SA-beads (2 μ1) for 30 min at +4°C. Samples were kept on ice and loaded one at a time on a BD LSREortessa^TM^ cell analyzer (BD Bioscience).

### Flow cytometry settings, data acquisition and analysis

Data acquisition was performed with a fluorescence-activated cell sorter (FACS) LSREortessa^TM^ flow cytometer (BD Biosciences) and the dedicated BD FACSDiva^TM^ software.

To excite and detect liposome-bound EGFP fluorescence emission (excitation/emission, 488/509) a 488 nm laser line together with a filter set of a 505 nm long-pass filter and a 530/30 nm filter was used. To detect Cy5 (excitation/emission, 496,565/670) 532 nm laser line together with a filter set of a 635 nm long-pass filter and a 670/30 nm filter was used.

Before any measurements were made, voltages in the photomultiplier tube (PMT) were adjusted accordingly to make streptavidin bead population fit into the linear range of the instrument as visually evaluated by scatter plot (FSC-A vs. SSC-A, Fig. 1c).

Subsequently, PMT was adjusted to accommodate both background fluorescence from the beads and sample fluorescence into the detection window. The typical count rate was below 200 events/second. Raw data was analyzed by using a non-commercial Flowing Software ver. 2.5 (Mr Perttu Terho; Turku Centre for Biotechnology, Finland; www.flowingsoftware.com), where the appropriate population of beads was gated and analyzed for their respective fluorescence intensities. Median fluorescence values were used for the subsequent data analysis as these are less sensitive for outliers than mean values.

**BTK-PH-EGFP K_D_ fitting**. To obtain minimal background, synthetic POPC was used in the K_D_ measurement instead of EggPC. Liposomes containing synthetic POPC lipids (80.5% w/w), cholesterol (15% w/w), biotinylated lipid (2% w/w) and PI(3,4,5)P_3_ (2.5% w/w) were prepared as before. In control liposomes, used to measure background fluorescence resulting from non-specific binding events, POPC concentration was increased (83% w/w) to compensate for the absence of
PI(3,4,5)P_3_. Cells expressing BTK-PH-EGFP were lysed, concentration of BTK-PH-EGFP determined by using equation (2) as described and serial dilutions of BTK-PH-EGFP (20-700 nM) were prepared. Binding of BTK-PH-EGFP to PI(3,4,5)P_3_-positive and control liposomes was measured by flow cytometry and background fluorescence was subtracted. The theoretical maximum fluorescence (Fmax) value was estimated by curve fitting:

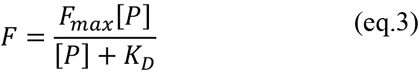

Where F is the raw background-subtracted fluorescence value and [P] is protein concentration. Raw fluorescence values were then normalized to F_max_ to determine occupancy:

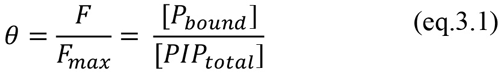

where [P_bound_] is the concentration of the protein bound to PIP and [PIP_total_] is the total concentration of PIP at the vesicle. Finally, K_D_ was calculated from equation:

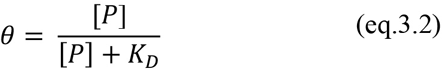

## Statistical analysis

Statistical significance was determined using the Student’s t-test (unpaired, two-tailed, unequal variance). N numbers are indicated in the figure legends. P-value of 0.05 was considered as a borderline for statistical significance.

## AUTHOR CONTRIBUTIONS

Conceptualization, N.dF. and J.I.; Methodology, N.dF., D.L., M.M.; Formal Analysis, N.dF. and M.M.; Investigation, N.dF., M.M., J.A., L.P. A.M.; Resources, B.G., P.M., B.T.G.; Writing-Original Draft, N.dF., H.H., J.I.; Visualization, N.dF. and H.H.; Supervision, J.I.; Funding Acquisition, J.I.

## ACKNOWLEDGMENTS

We thank P. Laasola, and J. Siivonen for excellent technical assistance. H. Bousquet and C. Kikuti (Institut Curie) are acknowledged for technical help. The CIC facilities at Turku Centre for Biotechnology are acknowledged for their help with flow cytometry and the PROTEX facility at the University of Leicester for generating the Talin FERM-EGFP domain construct. This study has been supported by the Academy of Finland and ERC Consolidator Grant 615258 (J.I). M.M. has been supported by the Drug Research Doctoral Program, University of Turku. The authors declare no conflict of interest.

## References

Ahram, M., Litou, Z. I., Fang, R. and Al-Tawallbeh, G. (2006). Estimation of membrane proteins in the human proteome. In. Silico Biol. 5, 379–386.

Almen, M. S., Nordstrom, K. J., Fredriksson, R. and Schioth, H. B. (2009). Mapping the human membrane proteome: a majority of the human membrane proteins can be classified according to function and evolutionary origin. BMC Biol., 50.

Ananthanarayanan, B., Stahelin, R. V., Digman, M. A. and Cho, W. (2003). Activation mechanisms of conventional protein kinase C isoforms are determined by the ligand affinity and conformational flexibility of their C1 domains. J. Biol. Chem. 47, 46886–46894.

Arinaminpathy, Y., Khurana, E., Engelman, D. M. and Gerstein, M. B. (2009). Computational analysis of membrane proteins: the largest class of drug targets. Drug Discov. Today 23–24, 11301135.

Besenicar, M., Macek, P., Lakey, J. H. and Anderluh, G. (2006). Surface plasmon resonance in protein-membrane interactions. Chem. Phys. Lipids 1-2, 169–178.

Bouvard, D., Pouwels, J., De Franceschi, N. and Ivaska, J. (2013). Integrin inactivators: balancing cellular functions in vitro and in vivo. Nat. Rev. Mol. Cell Biol. 7, 430–442.

Calderwood, D. A., Campbell, I. D. and Critchley, D. R. (2013). Talins and kindlins: partners in integrin-mediated adhesion. Nat. Rev. Mol. Cell Biol. 8, 503–517.

Ceccato, L., Chicanne, G., Nahoum, V., Pons, V., Payrastre, B., Gaits-Iacovoni, F. and Viaud, J. (2016). PLIF: A rapid, accurate method to detect and quantitatively assess protein-lipid interactions. Sci Signal 421, RS2.

Cullen, P. J. (2008). Endosomal sorting and signalling: an emerging role for sorting nexins. Nat. Rev. Mol. Cell Biol. 7, 574–582.

Elliott, P. R., Goult, B. T., Kopp, P. M., Bate, N., Grossmann, J. G., Roberts, G. C. K., Critchley, D. R. and Barsukov, I. L. (2010). The Structure of the talin head reveals a novel extended conformation of the FERM domain. Structure 10, 1289–1299.

Garcia, P., Gupta, R., Shah, S., Morris, A. J., Rudge, S. A., Scarlata, S., Petrova, V., McLaughlin, S. and Rebecchi, M. J. (1995). The pleckstrin homology domain of phospholipase C-delta 1 binds with high affinity to phosphatidylinositol 4,5-bisphosphate in bilayer membranes. Biochemistry 49, 1622816234.

Geertsma, E. R., Nik Mahmood, N a B, Schuurman-Wolters, G. K. and Poolman, B. (2008). Membrane reconstitution of ABC transporters and assays of translocator function. Nat Protoc 2, 256266.

Gillooly, D. J., Morrow, I. C., Lindsay, M., Gould, R., Bryant, N. J., Gaullier, J. M., Parton, R. G. and Stenmark, H. (2000). Localization of phosphatidylinositol 3-phosphate in yeast and mammalian cells. EMBO J. 17, 4577–4588.

Kojima, T., Fukuda, M., Watanabe, Y., Hamazato, F. and Mikoshiba, K. (1997). Characterization of the pleckstrin homology domain of Btk as an inositol polyphosphate and phosphoinositide binding domain. Biochem. Biophys. Res. Commun. 2, 333–339.

Kolena, J. (1989). Functional reconstitution of rat ovarian LH/hCG receptor into proteoliposomes. FEBS Lett. 2, 425–428.

Lacapère, J. J., Delavoie, F., Li, H., Pèranzi, G., Maccario, J., Papadopoulos, V. and Vidic, B. (2001). Structural and functional study of reconstituted peripheral benzodiazepine receptor. Biochem. Biophys. Res. Commun. 2, 536–541.

Lemmon, M. A. and Ferguson, K. M. (2000). Signal-dependent membrane targeting by pleckstrin homology (PH) domains. Biochem. J., 15;350 1–18.

Lemmon, M. A., Ferguson, K. M., O’Brien, R., Sigler, P. B. and Schlessinger, J. (1995). Specific and high-affinity binding of inositol phosphates to an isolated pleckstrin homology domain. Proc. Natl. Acad. Sci. U. S. A. 23, 10472–10476.

Lemmon, M. A. (2008). Membrane recognition by phospholipid-binding domains. Nat. Rev. Mol. Cell Biol. 2, 99–111.

Lévy, D., Bluzat, A., Seigneuret, M. and Rigaud, J. L. (1990). A systematic study of liposome and proteoliposome reconstitution involving Bio-Bead-mediated Triton X-100 removal. Biochim. Biophys. Acta 2, 179–190.

Moriyama, Y., Takano, T. and Ohkuma, S. (1984). Solubilization and reconstitution of a lysosomal H+-pump. J. Biochem. 3, 927–930.

Mouro-Chanteloup, I., Cochet, S., Chami, M., Genetet, S., Zidi-Yahiaoui, N., Engel, A., Colin, Y., Bertrand, O. and Ripoche, P. (2010). Functional reconstitution into liposomes of purified human RhCG ammonia channel. PLoS ONE 1, e8921.

Müller, D. J., Wu, N. and Palczewski, K. (2008). Vertebrate membrane proteins: structure, function, and insights from biophysical approaches. Pharmacol. Rev. 1, 43–78.

Nesper, J., Brosig, A., Ringler, P., Patel, G. J., Müller, S. A., Kleinschmidt, J. H., Boos, W., Diederichs, K. and Welte, W. (2008). Omp85(Tt) from Thermus thermophilus HB27: an ancestral type of the Omp85 protein family. J. Bacteriol. 13, 4568–4575.

Neves, P., Lopes Silvia, C D N, Sousa, I., Garcia, S., Eaton, P. and Gameiro, P. (2009). Characterization of membrane protein reconstitution in LUVs of different lipid composition by fluorescence anisotropy. J Pharm Biomed Anal 2, 276–281.

Pearce, L. R., Komander, D. and Alessi, D. R. (2010). The nuts and bolts of AGC protein kinases. Nat. Rev. Mol. Cell Biol. 1, 9–22.

Rameh, L. E. et al. (1997). A comparative analysis of the phosphoinositide binding specificity of pleckstrin homology domains. J. Biol. Chem. 35, 22059–22066.

Richard, P., Rigaud, J. L. and Gräber, P. (1990). Reconstitution of CF0F1 into liposomes using a new reconstitution procedure. Eur. J. Biochem. 3, 921–925.

Rigaud, J. L., Pitard, B. and Levy, D. (1995). Reconstitution of membrane proteins into liposomes: application to energy-transducing membrane proteins. Biochim. Biophys. Acta 3, 223–246.

Rossier, O. et al. (2012). Integrins β1 and β3 exhibit distinct dynamic nanoscale organizations inside focal adhesions. Nat. Cell Biol. 10, 1057–1067.

Saliba, A. et al. (2014). A quantitative liposome microarray to systematically characterize protein-lipid interactions. Nat. Methods 1, 47–50.

Smith, S. A. and Morrissey, J. H. (2004). Rapid and efficient incorporation of tissue factor into liposomes. J. Thromb. Haemost. 7, 1155–1162.

Song, A., Zhang, J., Zhang, M., Shen, T. and Tang, J. (2000). Spectral properties and structure of fluorescein and its alkyl derivatives in micelles. Colloids and Surfaces A: Physicochemical and Engineering Aspects 3, 253–262.

Stenmark, H., Aasland, R. and Driscoll, P. C. (2002). The phosphatidylinositol 3-phosphate-binding FYVE finger. FEBS Lett. 1, 77–84.

Temmerman, K. and Nickel, W. (2009). A novel flow cytometric assay to quantify interactions between proteins and membrane lipids. J. Lipid Res. 6, 1245–1254.

Winograd-Katz, S. E., Fässler, R., Geiger, B. and Legate, K. R. (2014). The integrin adhesome: from genes and proteins to human disease. Nat. Rev. Mol. Cell Biol. 4, 273–288.

Worrall, J. A. R. and Mason, J. M. (2011). Thermodynamic analysis of Jun-Fos coiled coil peptide antagonists. FEBS J. 4, 663–672.

Wu, H., Henzie, J., Lin, W., Rhodes, C., Li, Z., Sartorel, E., Thorner, J., Yang, P. and Groves, J. T. (2012). Membrane-protein binding measured with solution-phase plasmonic nanocube sensors. Nat. Methods 12, 1189–1191.

Ye, F., Hu, G., Taylor, D., Ratnikov, B., Bobkov, A. A., McLean, M. A., Sligar, S. G., Taylor, K. A. and Ginsberg, M. H. (2010). Recreation of the terminal events in physiological integrin activation. J. Cell Biol. 1, 157–173.

Young, H. S., Rigaud, J. L., Lacapère, J. J., Reddy, L. G. and Stokes, D. L. (1997). How to make tubular crystals by reconstitution of detergent-solubilized Ca2(+)-ATPase. Biophys. J. 6, 2545–2558.

Zhang, X., Zhang, J. and Liu, L. (2014). Fluorescence properties of twenty fluorescein derivatives: lifetime, quantum yield, absorption and emission spectra. J Fluoresc 3, 819–826.

Zhao, H. and Lappalainen, P. (2012). A simple guide to biochemical approaches for analyzing protein-lipid interactions. Mol. Biol. Cell 15, 2823–2830.

